# A genome-wide screen in macrophages defines host genes regulating the uptake of *Mycobacterium abscessus*

**DOI:** 10.1101/2022.12.20.521338

**Authors:** Haleigh N. Gilliland, Olivia K. Beckman, Andrew J. Olive

## Abstract

The interactions between a host cell and a pathogen can dictate disease outcomes and are important targets for host-directed therapies. *Mycobacterium abscessus* (Mab) is a highly antibiotic resistant, rapidly growing non-tuberculous mycobacterium that infects patients with chronic lung diseases. Mab can infect host immune cells, such as macrophages, which contribute to its pathogenesis. However, our understanding of initial host-Mab interactions remains unclear. Here, we developed a functional genetic approach to define these host-Mab interactions by coupling a Mab fluorescent reporter with a genome-wide knockout library in murine macrophages. We used this approach to conduct a forward genetic screen to define host genes that contribute to the uptake of Mab by macrophages. We identified known regulators of phagocytosis, such as the integrin ITGB2, and uncovered a key requirement for glycosaminoglycan (sGAG) synthesis for macrophages to efficiently take up Mab. CRISPR-Cas9 targeting of three key sGAG biosynthesis regulators, *Ugdh, B3gat3 and B4galt7* resulted in reduced uptake of both smooth and rough Mab variants by macrophages. Mechanistic studies suggest that sGAGs function upstream of pathogen engulfment and are required for the uptake of Mab, but not *Escherichia coli* or latex beads. Further investigation found that the loss of sGAGs reduced the surface expression, but not the mRNA expression, of key integrins suggesting an important role for sGAGs in modulating surface receptor availability. Together, these studies globally define and characterize important regulators of macrophage-Mab interactions and are a first step to understanding host genes that contribute to Mab pathogenesis and disease.

**IMPORTANCE:** Pathogen interactions with immune cells like macrophages contribute to pathogenesis, yet the mechanisms underlying these interactions remain largely undefined. For emerging respiratory pathogens, like *Mycobacterium abscessus*, understanding these host-pathogen interactions is important to fully understand disease progression. Given that *M. abscessus* is broadly recalcitrant to antibiotic treatments, new therapeutic approaches are needed. Here, we leveraged a genome-wide knockout library in murine macrophages to globally define host genes required for *M. abscessus* uptake. We identified new macrophage uptake regulators during *M. abscessus* infection, including a subset of integrins and the glycosaminoglycan synthesis (sGAG) pathway. While ionic characteristics of sGAGs are known to drive pathogen-cell interactions, we discovered a previously unrecognized requirement for sGAGs to maintain robust surface expression of key uptake receptors. Thus, we developed a flexible forward-genetic pipeline to define important interactions during *M. abscessus* infection and more broadly identified a new mechanism by which sGAGs control pathogen uptake.

## INTRODUCTION

*Mycobacterium abscessus* (Mab) is a rapidly growing non-tuberculous mycobacterium (NTM) that causes opportunistic infections in patients with chronic lung diseases, like cystic fibrosis and chronic obstructive pulmonary disease (COPD) (1, 2). Mab is the second most common NTM respiratory pathogen recovered in the United States, accounting for a significant number of rapidly growing mycobacterial respiratory disease isolates (3). Due to its recalcitrance to many antibiotics, current treatment success rates remain below 50% (4-6). Treatment is further complicated by the ability of Mab to transition from a smooth to rough morphology that drives biofilm formation and decreases antibiotic sensitivity (7). Thus, Mab is an emerging pathogen of clinical importance and there is a critical need to develop new treatment options.

Developing more effective therapies requires a deeper understanding of Mab pathogenesis and host-pathogen interactions that drive infection. Recent work suggests that Mab interactions with macrophages are critical for disease progression (3, 8, 9). Both smooth and rough Mab variants can survive in macrophages with rough variants initiating more rapid cell death cascades (3, 8, 9). Transposon mutagenesis studies and other genetic approaches in Mab have identified many genes that are required for effective antibiotic killing and intracellular survival, including the ESX-IV system (10-12). This virulence determinant contributes to the inhibition of lysosomal fusion enabling intracellular survival of Mab in macrophages (12). In contrast to what is known about Mab, little is known about macrophage genes required for Mab uptake and survival. While studies have shown that Mab interacts with TLR2 and Dectin-1 to initiate uptake, a systematic characterization of Mab uptake by macrophages remains lacking (13, 14).

Uptake of pathogens requires effective interactions between the pathogen and macrophages. Macrophages use surface pattern recognition receptors (PRRs), including toll-like receptors, mannose receptors and scavenger receptors, for the initial recognition of many pathogens (15-19). Other modifications to surface proteins, including sulfated glycosaminoglycan (sGAG) modifications such as heparin sulfate and chondroitin sulfate, affect the efficiency of pathogen attachment to the cell surface (20, 21). On epithelial cells, sGAGs play an important role for the attachment of several pathogens including Coronavirus and *Chlamydia trachomatis* (22-24). It is predicted that these surface modifications alter the charge interactions between pathogens and the host cell surface to modulated initial attachment (25, 26).

Following adherence, phagocytosis or receptor mediated engulfment of pathogens enables internalization. While each phagocytic cargo has unique components, the general pathway for phagocytosis is conserved. Phagocytosis begins with the generation of a phagosome following a ligand binding to macrophage surface receptors (27). This binding event activates downstream signaling cascades that remodel the actin cytoskeleton to drive the progressive engagement of additional receptors around the particle (28, 29). After engulfment, controlled membrane fusion events result in the formation of the nascent phagosome. The phagosome then interacts with various endosome components that acidify the compartment and ultimately lead to compartment fusion with the lysosome (30-35). Several elegant studies have dissected unique pathways that contribute to phagocytosis of distinct cargos, defining receptors and signaling cascades that are essential (36-38). However, no studies have globally examined host genes that control Mab interactions with macrophages.

Here, we defined the host genes that control the uptake of Mab into macrophages by coupling a brightly fluorescent Mab reporter with a genone-wide CRISPR-Cas9-mediated macrophage knockout library in murine immortalized bone marrow-derived macrophages (iBMDMs). We performed a forward genetic screen to enrich for macrophages that could or could not take up Mab identifying several previously defined phagocytosis components. Follow up studies uncovered a strong requirement for sGAG production in macrophages for efficient Mab uptake. While loss of sGAGs did not affect general phagocytosis pathways or opsonized uptake of Mab, sGAGs were essential for initial interactions of both smooth and rough Mab variants with macrophages. Mechanistic studies uncovered a role for sGAGs in maintaining high surface expression of the key integrins ITGB2 and ITGAL, suggesting a new role for sGAGs in maintaining receptor availability on the cell surface. These results uncover important host pathways that modulate the interactions between Mab and macrophages during infection.

## RESULTS

### Fluorescent Mab reporter enables dissection of Macrophage-Mab interactions

While Mab infects macrophages, our understanding of these interactions remains limited. To help address this gap in knowledge we developed a brightly fluorescent Mab reporter strain. A constitutive mEmerald GFP vector was transformed into the smooth variant of Mab strain ATCC-19977. Comparisons to non-fluorescent Mab showed that bacterial growth and antibiotic mediated killing with rifampicin in liquid media were unaffected by fluorescent protein expression (Figure 1A). We next examined the utility of the fluorescent Mab strain to dissect host-pathogen interactions with macrophages. We first examined the robustness of the fluorescent reporter by infecting immortalized bone marrow derived macrophages (iBMDMs) from C57BL/6J mice. One previous study using a GFP fluorescent Mab was unable to detect increased fluorescence in macrophages until 24 hours of infection, when attachment, uptake and bacterial growth all contribute to differential fluorescence (39). We hypothesized that earlier timepoints could be examined using the brighter mEmerald GFP, allowing us to distinguish uptake from intracellular growth. To test this hypothesis, we quantified the number of mEmerald positive macrophages by flow cytometry six hours after infecting iBMDMs with fluorescent Mab at increasing multiplicities of infection (MOIs). We noted that increasing the MOI resulted in an increase in the percent of mEmerald+ cells and the mean fluorescence intensity of these infected cells (Figure 1B-1D). We next quantified the mEmerald+ cells over the first six hours of infection of iBMDMs at an MOI of 5. We observed the number of infected cells 2-hours post-infection showed a minimal increase compared to uninfected controls, while there was a significant increase in infected cells at four- and six-hours post-infection (Figure 1E and 1F). Thus, the mEmerald GFP reporter does not affect Mab growth while enabling the dissection of early interactions between Mab and macrophages.

**Figure 1.**
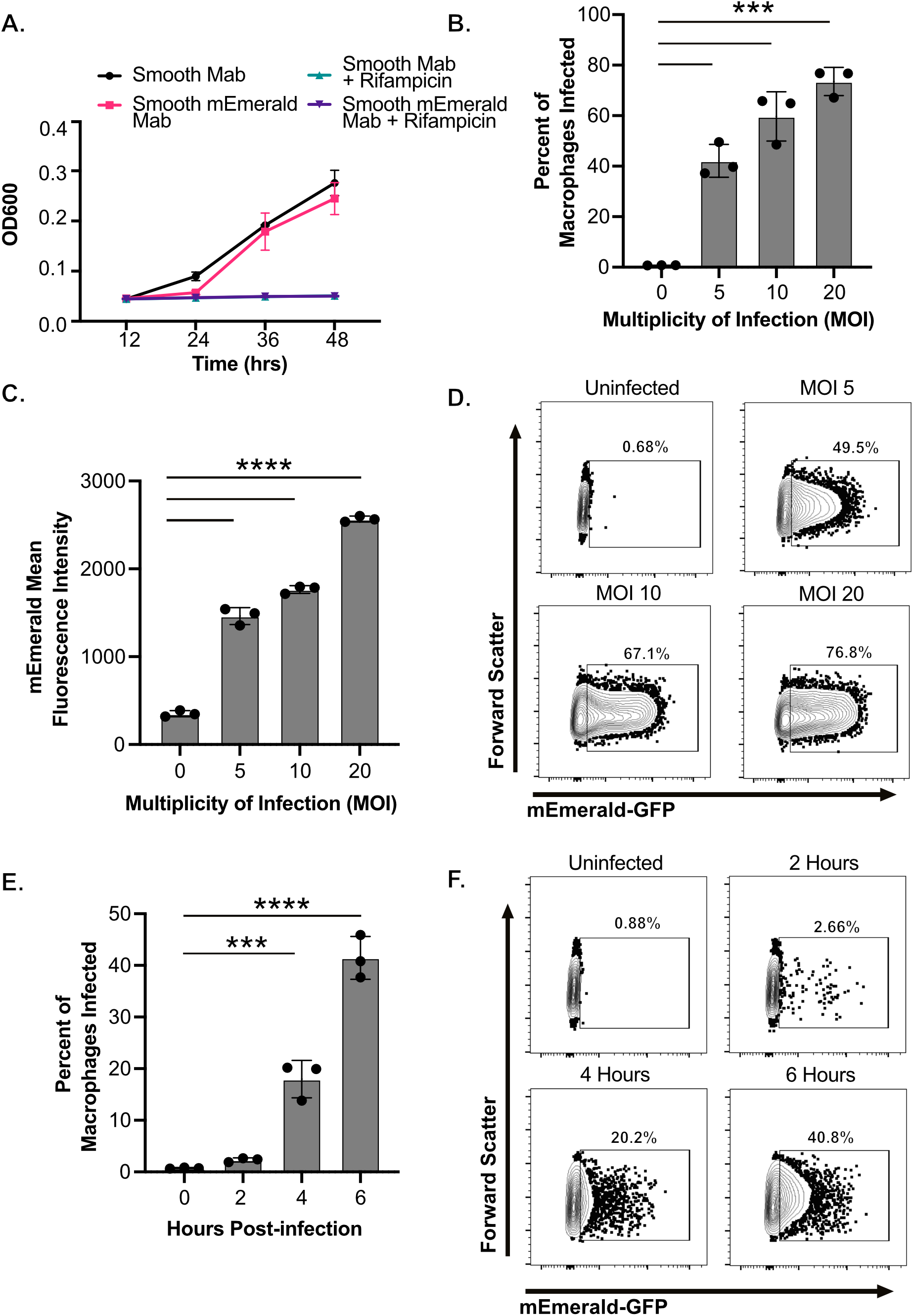
Early interactions with fluorescent *Mycobacterium abscessus* and macrophages can be detected by flow cytometry. **(A)** The optical density of control or mEmerald transformed *M. abscessus* ATCC-19977 was monitored over 48 hours in 7H9 broth in the presence or absence of Rifampicin (128 μg/ml). **(B-D)** iBMDMs from C57BL6J mice were infected with increasing MOI (5-20) of mEmerald *M. abscessus* for six hours. Flow cytometry was used to measure (B) the percent of cells that were infected (mEmerald+) and (C) the mean fluorescence intensity of infected cells. (D) Representative flow cytometry plots gated on live and single cells are shown for each MOI. Data are from one of three independent experiments. **(E-F)** iBMDMs from C57BL6/J mice were infected with mEmerald *M. abscessus* at an MOI of 5 for two, four or six hours. **(E)** Flow cytometry was used to measure the percent of cells that were infected (mEmerald+). **(F)** Shown are representative flow cytometry plots gated on live and single cells for each time point. All results shown are representative results from one of three independent experiments. ****p<.0001 ***p<.001 by one-way ANOVA with Dunnett’s multiple comparison test.

### CRISPR-Cas9 loss-of-function (LOF) screen identifies host genes required for Mab uptake by macrophages

Host pathways required for macrophage uptake of Mab remain almost entirely unknown. To test if the flow cytometry-based uptake assay could identify host pathways required for Mab uptake, we used a known phagocytosis inhibitor, Cytochalasin D, to inhibit actin polymerization (40). iBMDMs were pretreated with Cytochalasin D then infected with mEmerald Mab. The percent of cells that were infected were then quantified four hours later by flow cytometry. We observed a nearly 50% decrease in mEmerald+ macrophages treated with Cytochalasin D compared to vehicle controls (Figure 2A and 2B). These data highlight the sensitivity of the uptake assay to dissect early interactions between Mab and macrophages.

**Figure 2.**
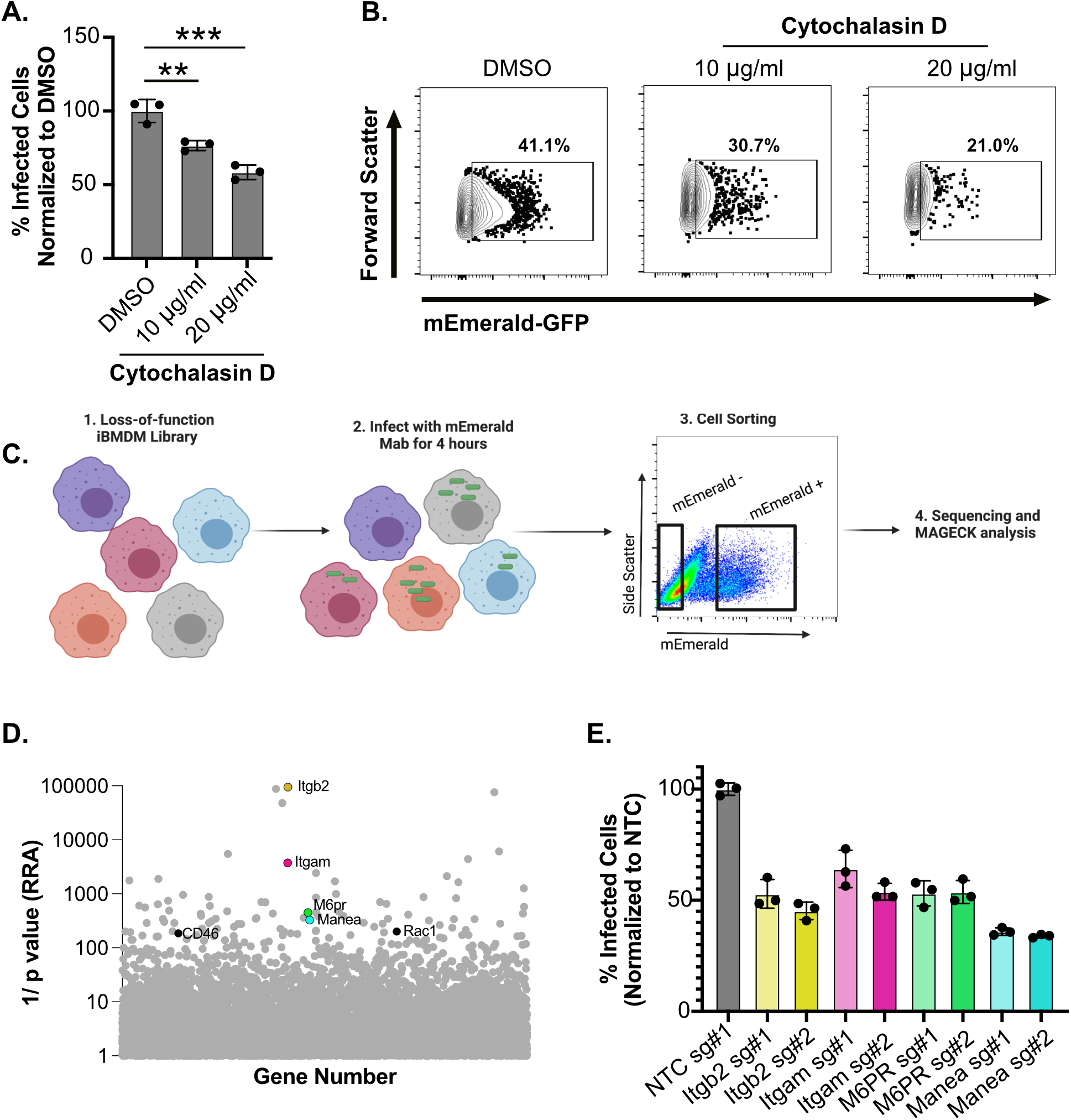
Genome-wide loss-of-function screen in iBMDMs identifies host genes required for uptake of *M. abscessus*. **(A)**. iBMDMs were treated with DMSO or Cytochalasin D (10μg/ml or 20μg/ml) for two hours then infected with mEmerald *M. abscessus* at an MOI of 5 for four hours. The percent uptake was quantified by flow cytometry. Results are normalized to the mean percent uptake of the NTC+DMSO condition **(B)** Shown are representative flow cytometry plots gated on live single cells for each treatment. **(C)** A schematic of the genome-wide screen to identify host genes that are required for *M. abscessus* uptake by macrophages. A genome-wide CRISPR-Cas9 library generated in Cas9+ iBMDMs with sgRNAs from the Brie library (4 sgRNAs per gene) was infected with mEmerald *M. abscessus* for four hours and mEmerald^+^ and mEmerald^-^ populations were isolated by FACS. The representation of sgRNAs in each population were determined by sequencing. Figure made in Biorender. **(D)** Shown is the score determined by the alpha-robust rank algorithm (α-RRA) in MAGeCK for each gene in the CRISPR-Cas9 library that passed filtering metrics from three independent screen replicates. Highlighted genes represent known host factors that contribute to uptake of pathogens. **(E)** Cas9+ iBMDMs targeted with the indicated sgRNAs for candidates (2 per candidate gene) were infected with mEmerald *M. abscessus* for 4 hours at an MOI of 5. The uptake of *M. abscessus* was quantified by flow cytometry. These data are normalized to the mean percent of uptake of *M. abscessus* by non-targeting control cells. The results are representative of at least three independent experiments. ***p<.001 **p<.01 by one-way ANOVA with Dunnett’s multiple comparison test. For the data in panel 2E each candidate sgRNA is p<.001 compared to the NTC.

Next, we leveraged the uptake assay to globally identify host regulators required for Mab uptake by macrophages (Figure 2C). We previously generated a robust and reproducible pooled genome-wide loss-of-function library in Cas9+ iBMDMs (41). Knockouts in this pool were generated with sgRNAs from the Brie library which targets four independent single guide RNA molecules (sgRNAs) per coding gene and over 1,000 non-targeting controls (NTC) (42). To identify regulators of Mab uptake, we infected the loss-of-function library of macrophages with mEmerald Mab for four hours. Fluorescence activated cells sorting (FACS) was then used to isolate over 200x sgRNA coverage of the library from mEmerald+ cells and mEmerald-cells. Following genomic DNA extraction, sgRNA abundances for each sorted bin were quantified by deep sequencing.

To test for statistical enrichment of sgRNAs and genes, we used the modified robust rank algorithm (α-RRA) employed by Model-based Analysis of Genome-wide CRISPR/Cas9 Knockout (MAGeCK). This algorithm ranks sgRNAs by effect before filtering low ranking sgRNAs to improve significance testing. To identify macrophage genes required for Mab uptake during early infection, we compared the enrichment of sgRNAs in the mEmerald -(Mab uninfected) directly to the mEmerald+ (Mab infected) population (Table S1). The α-RRA analysis identified 100 genes with a p value <0.01 and a fold change of 2 with at least 2 independent sgRNAs. Among the top 100 candidates, we identified known regulators of macrophage phagocytosis including *Manea, M6PR, Itgb2* (*CD18*), *Itgam* (*CD11b*), *CD46* and *Rac1* with each gene showing enrichment in the mEmerald-population (Figure 2D) (43). Guide-level analysis showed agreement with all four sgRNAs targeting these genes suggesting they are bona fide hits and that our screen was robust (Table S1).

To confirm the screen results, we validated the role of *Manea, M6PR, Itgb2* and *Itgam* in controlling Mab uptake by macrophages. We generated two independent iBMDM lines targeting two distinct sgRNAs per gene, in addition to a non-targeting control. These cells were then infected with mEmerald Mab and uptake was quantified four hours later. In agreement with our screen results, we found that targeting each candidate gene resulted in a significant decrease in mEmerald+ iBMDMs compared to NTC cells (Figure 2E). Taken together these results show that the genome-wide screen identified host genes that contribute to early interactions between Mab and macrophages.

### Sulfated glycosaminoglycans (sGAGs) are required for macrophage uptake of Mab

To discover new pathways that are required for Mab uptake by macrophages, we next used DAVID analysis and gene set enrichment analysis (GSEA) to identify functional enrichments from our uptake screen dataset (Table S2). These analyses identified the sulfated glycosaminoglycan (sGAG) synthesis pathway as required for efficient uptake of Mab (Figure 3A and Table S2). While a variety of pathogens use sGAGs to facilitate attachment and invasion of epithelial cells, little is known about their role in macrophages. Within the top candidates in the screen were multiple genes within the sGAG biosynthesis pathway (Figure 3B – Blue Dots). Three of these genes, *UGDH, B3GAT3*, and *B4GALT7* encode UDP-glucose 6-dehydrogenase, galactosyltransferase I and glucuronosyltransferase I, respectively and were among the top 30 candidates while showing strong agreement among the four independent sgRNAs (Table S1). These data suggest that sGAGs contribute to Mab uptake by macrophages.

**Figure 3.**
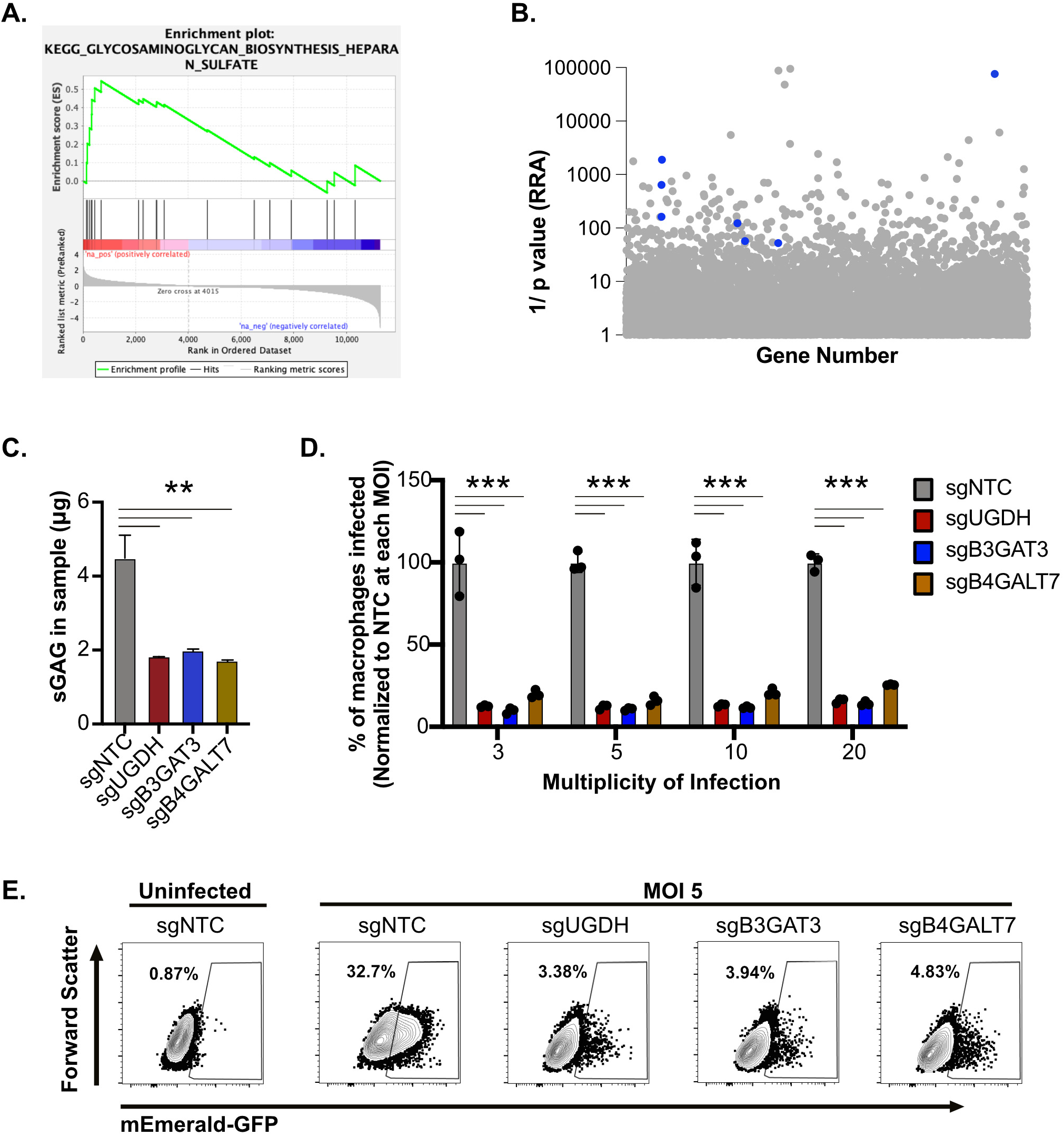
Sulfated glycosaminoglycans are required for effective uptake of *M. abscessus* by macrophages. **(A)** Gene set enrichment analysis was used to identify enriched pathways from the ranked forward genetic screen. Shown is a leading-edge analysis plot representing a top enriched KEGG pathway related to glycosaminoglycan biosynthesis. **(B)** The α-RRA ranking of sGAG biosynthesis related genes identified in the genome-wide screen highlighted as blue dots. **(C)** Total sGAGs were quantified in sgNTC, sgUGDH, sgB3GAT3 and sgB4GALT7 iBMDMs and normalized to total protein content. Shown is a representative result from four independent experiments. **(D)** sgNTC, sgUGDH, sgB3GAT3 and sgB4GALT7 iBMDMs were infected with mEmerald *M. abscessus* at increasing MOI for four hours then the percent infected cells were quantified by flow cytometry. Shown is the percent of infected cells normalized to the mean of the uptake of NTC at each MOI. These results are representative of four independent experiments. **(E)** Shown is a representative flow cytometry plot gated on live and single cells for the indicated genotypes at an MOI of 5. ***p<.001 **p<.01 by one-way ANOVA with Dunnett’s multiple comparison test.

To validate the importance of macrophage sGAGs for Mab uptake, we targeted *Ugdh, B3gat3* and *B4galt7* directly in iBMDMs. Using an approach targeting two independent sgRNAs simultaneously we generated a panel of iBMDMs with editing efficiency ranging from 60-95% in each gene, and one cell line per gene was selected for follow up studies (42). We first examined functional differences following gene editing by quantifying the total sGAGs in sGAG-targeted and NTC iBMDMs. We found that iBMDMs targeted for *Ugdh, B3gat3* and *B4galt7*each resulted in a significant reduction in sGAGs compared to NTC macrophages (Figure 3C). We next used these sGAG-targeted cells to examine differences in Mab uptake. NTC and sGAG-targeted iBMDMs were infected with mEmerald Mab for four hours with increasing MOIs and bacterial uptake was quantified. At each MOI we observed a significant decrease in Mab uptake in sGAG-targeted iBMDMs compared to the NTC cells (Figure 3D and 3E). These results confirm that sGAGs contribute to the effective uptake of Mab by macrophages, validating our genome-wide screen and bioinformatic analysis.

### sGAGs are required for macrophage uptake of rough Mab variants

While our data suggest sGAGs are needed for uptake of smooth Mab variants by macrophages, it remained unclear if this host pathway was also required for rough variant uptake. To directly test this question, a rough Mab variant derived from ATCC 19977 was transformed with the mEmerald reporter. Similar to the smooth variant, we noted no defects in growth or antibiotic killing in broth culture (Figure 4A) and robust uptake by iBMDMs in an MOI dependent manner (Figure 4B-4D). To test if sGAGs are also required for uptake of rough Mab we next infected NTC and sGAG-targeted iBMDMs with the rough mEmerald Mab reporter for four hours at increasing MOI and quantified the percentage of infected cells. We observed over a 50% decrease of rough Mab uptake in sGAG-targeted macrophages compared to control macrophages at all MOIs (Figure 4E and 4F). Thus, sGAGs are required for macrophage uptake of both smooth and rough Mab variants.

**Figure 4.**
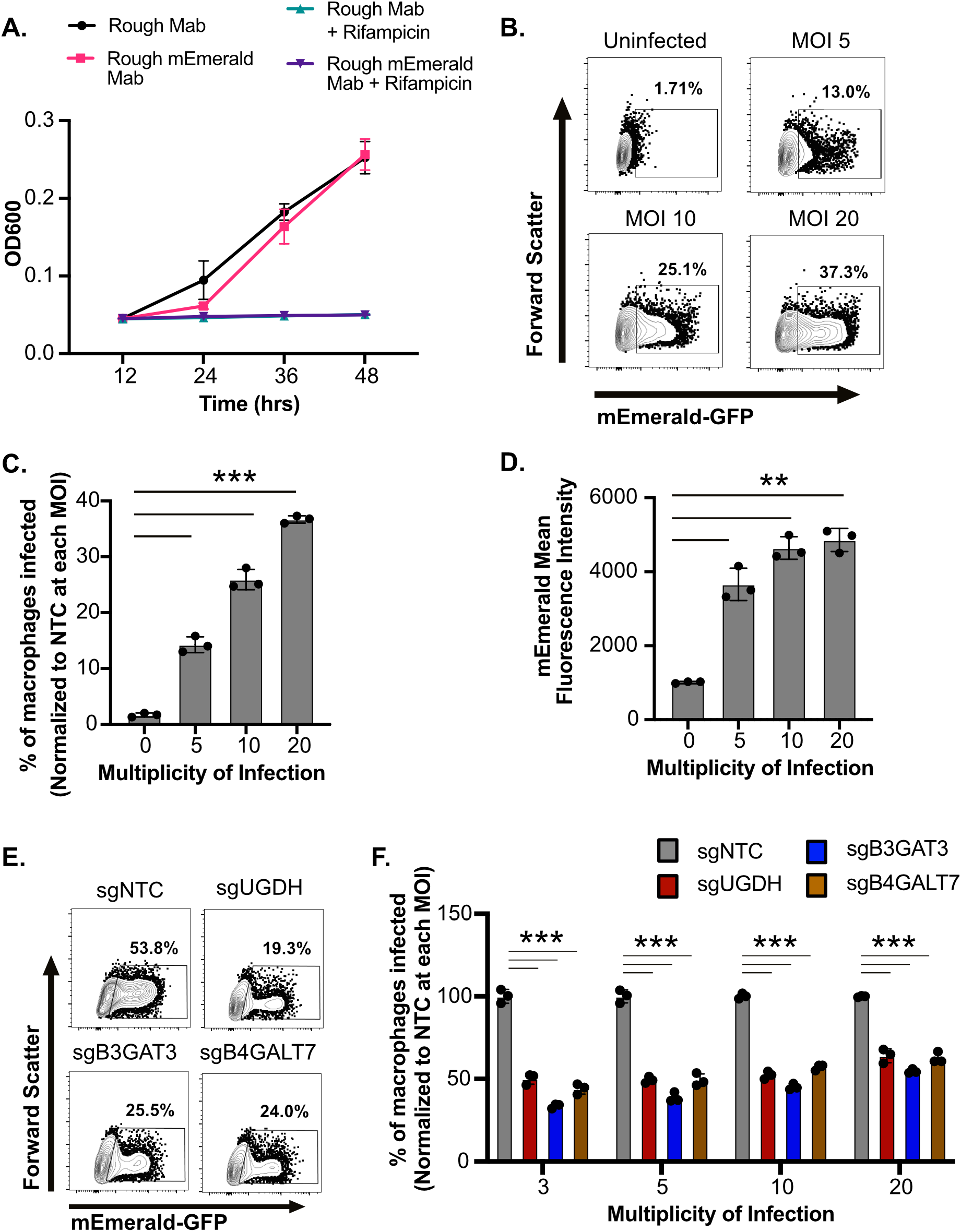
Rough variants of *M. abscessus* require sGAGs for efficient uptake by macrophages. **(A)** The optical density of control or mEmerald transformed rough variant of *M. abscessus* ATCC-19977 was monitored over 48 hours in 7H9 broth in the presence or absence of Rifampicin (128 μg/ml). **(B)** iBMDMs from C57BL6/J mice were infected with increasing MOI (5-20) of mEmerald rough variant of *M. abscessus* for four hours and flow cytometry was used to measure the percent of cells that were infected (mEmerald+). Shown are representative flow cytometry plots gated on live and single cell for an MOI of five. **(C)** The percent of macrophages infected with rough Mab and **(D)** the mean fluorescence intensity of infected cells at the indicated MOIs. **(E)** sgNTC, sgUGDH, sgB3GAT3 and sgB4GALT7 iBMDMs were infected with mEmerald rough variant of *M. abscessus* at increasing MOIs for four hours then the percent uptake of cells was quantified by flow cytometry. Shown is a representative flow cytometry plot gated on live and single cells for the indicated genotypes infected at an MOI of 5. **(F)** Shown is the percent of infected cells normalized to the mean of the percent uptake for the NTC at each MOI. Results in (A-F) are all representative of at least three independent experiments. ***p<.001 **p<.01 by one-way ANOVA with Dunnett’s multiple comparison test.

### sGAGs function upstream of Mab internalization

We next examined the mechanisms underlying sGAG-mediated control of Mab uptake. To understand if sGAGs control specific or general uptake pathways, we tested whether sGAGs were necessary for uptake of other phagocytic cargo by macrophages. First, we quantified the uptake of latex beads by incubating yellow-green-labeled beads at increasing concentrations with control or sGAG-targeted iBMDMs. In contrast to our results with Mab, we found no differences in the uptake of latex beads between control or sGAG-targeted iBMDMs (Figure 5A and 5B). We next examined whether sGAGs contribute to the uptake of *Escherichia coli*, a gram-negative bacterium. mEmerald expressing *E. coli* was incubated with NTC or sGAG-targeted iBMDMs for four hours then uptake was quantified. Similar to latex beads, we observed no significant differences in *E. coli* uptake between control or sGAG-targeted macrophages (Figure 5C and 5D). Taken together, these results suggest that sGAG-mediated uptake in macrophages occurs independently of general phagocytosis mechanisms and has pathogen specificity.

**Figure 5.**
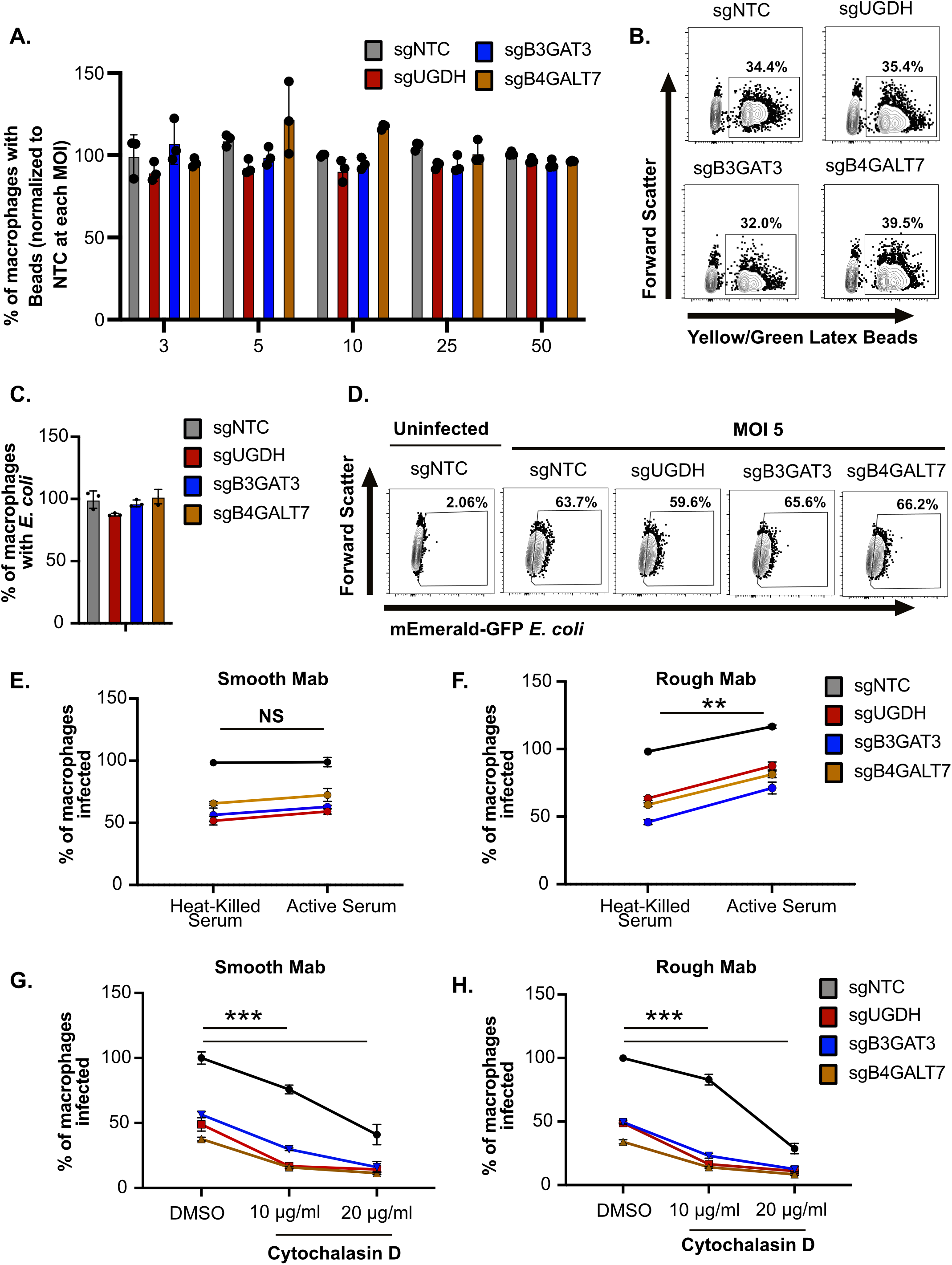
sGAGs do not alter general macrophage phagocytosis mechanisms and require viable bacteria to contribute to uptake. **(A)** Yellow-green-labeled latex beads were added at increasing concentrations (Bead:Cell ratio 3-50) to sgNTC, sgUGDH, sgB3GAT3 and sgB4GALT7 iBMDMs for four hours then the percent of cells with beads was quantified by flow cytometry. Shown is the percent of cells with beads normalized to the mean of the percent uptake of NTC at each concentration. **(B)** Shown are representative flow cytometry plots gated on live and single cells for the indicated genotypes treated with a bead:cell ratio of 5. **(C)** The indicated genotypes of iBMDMs were infected with mEmerald *E. coli* for four hours at an MOI of five. The percent infected cells were quantified by flow cytometry and were normalized to the mean of the percent uptake of NTC. **(D)** Shown are representative flow cytometry plots gated on live and single cells for the indicated genotyped infected with E. coli at an MOI of 5 **(E)** Smooth or **(F)** rough variants of mEmerald *M. abscessus* were incubated with heat-killed or active fetal bovine serum for thirty minutes then used to infect iBMDMs of the indicated genotypes for four hours at an MOI of five. The percent infected cells were quantified by flow cytometry and were normalized to the mean of sgNTC uptake in heat-killed serum for either smooth or rough variants. **(G)** iBMDMs of the indicated genotypes were treated with DMSO or Cytochalasin D (10μg/ml or 20μg/ml) for two hours then infected with mEmerald *M. abscessus* smooth or **(H)** rough variants at an MOI of 5 for four hours. The percent uptake was quantified by flow cytometry and all samples were normalized to the mean of sgNTC uptake in DMSO for either smooth or rough variants. All results are representative of at least 3 independent experiments with similar results. ***p<.001 **p<.01 by two-way ANOVA with Dunnett’s multiple comparison test.

Given that the uptake of pathogens can be influenced by opsonization, we next examined whether the sGAG pathway overlaps with complement-mediated uptake mechanisms. Both smooth and rough mEmerald Mab reporters were incubated in active or heat killed serum for thirty minutes. Following serum incubation, control or sGAG-targeted iBMDMs were infected and the percent uptake of Mab was quantified four hours later. Opsonization of the smooth Mab variant with active serum resulted in no significant change in uptake compared to heat-inactivated serum (Figure 5E). For the rough Mab variant, while uptake was significantly increased following incubation with active serum in all cells, the uptake differences between NTC and sGAG-targeted iBMDMs remained significant (Figure 5F). These data show that while smooth and rough Mab variants are differentially susceptible to complement-mediated phagocytosis, this uptake pathway is independent of sGAGs.

To distinguish if sGAGs are required for internalization or attachment of Mab we next treated control or sGAG-targeted macrophages with Cytochalasin D. We hypothesized that if sGAGs control initial attachment that blocking actin polymerization with Cytochalasin D would further inhibit Mab uptake in sGAG-targeted macrophages. To test this hypothesis, we pre-treated NTC and sGAG-targeted macrophages with DMSO or Cytochalasin D for two hours. Cells were then infected with smooth or rough variants of mEmerald Mab and uptake was quantified four hours later. We found that Cytochalasin D treatment significantly reduced uptake of both smooth and rough Mab variants compared to vehicle controls in all cell lines tested (Figure 5G and 5H). These data suggest that sGAGs are needed upstream of bacterial internalization as they function additively with actin polymerization inhibitors.

### Loss of sGAGs reduces the surface integrin expression on macrophages

While sGAGs directly modulate ionic interactions at the cellular surface, it remained possible that sGAG-modifications also regulate the surface levels of the receptors that are required for pathogen uptake (26). Given the importance of integrins for Mab uptake from our screen, we examined the expression of a subset of integrins including ITGB2, ITGAM, and ITGAL. When we quantified the mRNA levels of these integrins we found no changes in the mRNA expression of integrins between NTC and sGAG-targeted macrophages (Figure 6A). In contrast, while we observed no difference in the surface expression of ITGAM, we observed a significant decrease in ITGB2 and ITGAL expression on the surface of sGAG-targeted iBMDMs (Figure 6B and 6C). We next directly examined surface levels of the ITGB2/ITGAL heterodimer (LFA-1) and found significantly reduced expression on the surface of sGAG-targeted iBMDMs (Figure 6D and 6E). Taken together these data suggest that sGAGs modulate the surface expression of key integrins that contribute to Mab uptake by macrophages.

**Figure 6.**
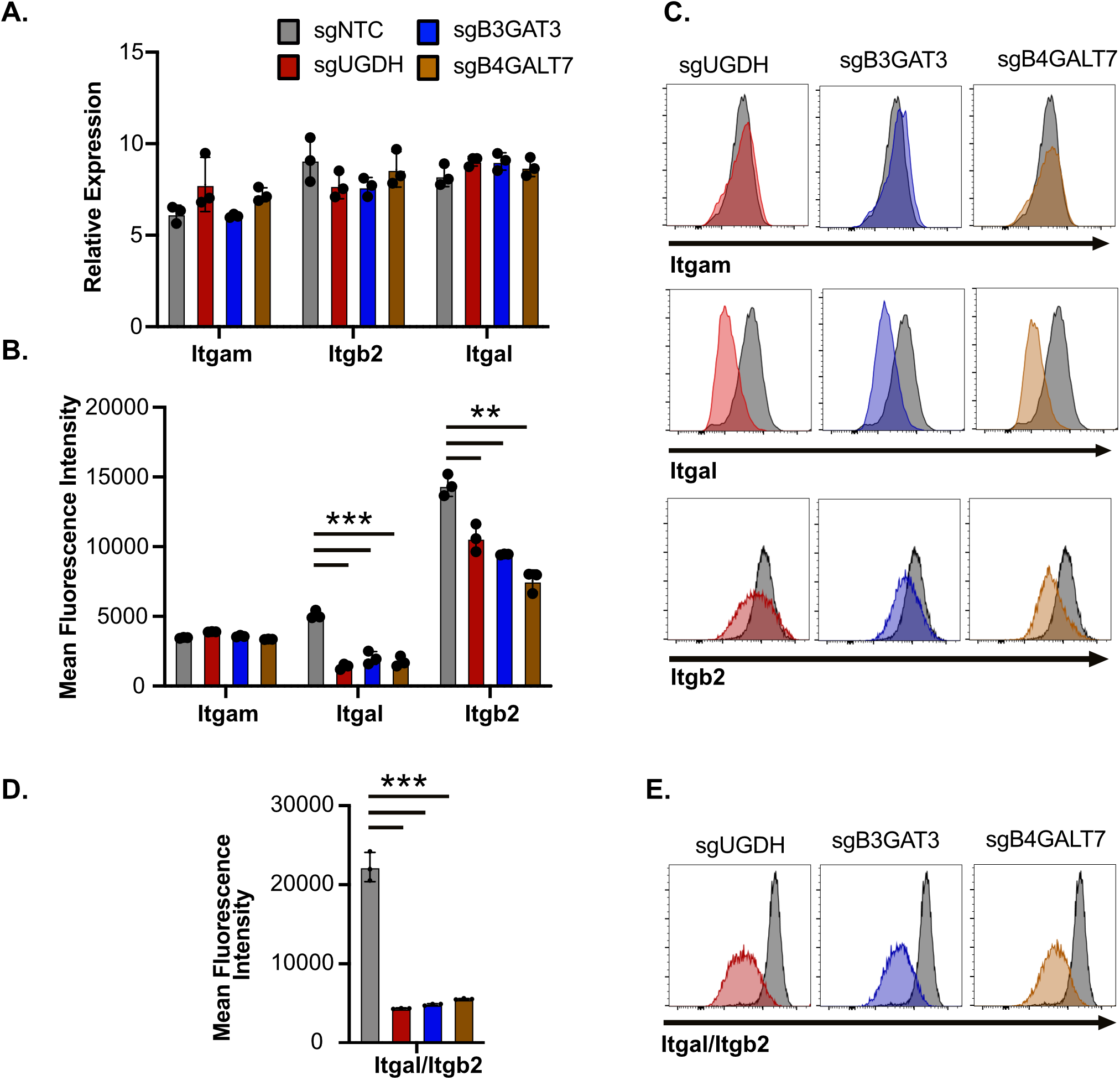
Reduced sGAGs results in lower surface expression of integrins on macrophages. **(A)** RNA was isolated from sgNTC, sgUGDH, sgB3GAT3 and sgB4GALT7 iBMDMs and the expression of *Itgb2, Itgal* and *Itgam* relative to *Gapdh* was quantified by qRT-PCR. (B) The surface expression of ITGB2, ITGAL and ITGAM was quantified in NTC and sGAG-targeted iBMDMs by flow cytometry. Shown is the mean fluorescence intensity of each integrin and (C) a representative histogram for each sGAG-targeted gene for each integrin overlaid with NTC is shown. (D) The surface expression of ITGB2-ITGAL heterodimers (LFA-1) were quantified on NTC and sGAG-targeted iBMDMs. Shown is the quantification of the mean fluorescence intensity and (F) a representative histogram for each iBMDM genotype overlaid with NTC. All results are representative of at least 2-3 independent experiments with similar results. ***p<.001 **p<.01 by one-way ANOVA with Dunnett’s multiple comparison test.

## DISCUSSION

Infections with *M. abscessus* are on the rise globally and their recalcitrance to antibiotic therapy is driving treatment failure (44-46). Designing effective treatments against Mab, including host-directed therapies, will require a fundamental understanding of critical host-Mab interactions that occur during infection. Here, we developed a Mab fluorescent reporter-based approach to investigate interactions of Mab with macrophages and conducted a forward genetic screen to globally identify host genes that are required for Mab uptake. Our screen identified many candidates including previously described players in pathogen uptake such as *Itgb2, Itgam, Manea* and *M6pr* (47, 48) which together with our validation highlight the robustness of this dataset and the strengths of the functional genetic approach.

As part of our bioinformatic analysis to uncover new pathways required for efficient Mab uptake, we identified a strong signature for the sGAG biosynthesis pathway. sGAGs are modified glycosaminoglycans, which consist of repeating polysaccharides attached to the cell surface and surface proteins (26). We targeted three independent genes in the sGAG synthesis pathway that all resulted in decreased uptake of both smooth and rough Mab variants by macrophages. While sGAGs have previously been shown to modulate host-pathogen interactions in epithelial cells, very little is known about their role in macrophages (22-24). We found that sGAGs did not affect latex bead or *E. coli* uptake, showing that general macrophage uptake mechanisms are intact in sGAG-targeted cells. In addition, Cytochalasin D treatment of sGAG-targeted macrophages further reduced Mab uptake, suggesting sGAGs contribute upstream of the actin polymerization required for Mab internalization. Together these data point towards a role for sGAGs in the adherence of Mab to macrophages. While many models suggest sGAGs modulate ionic interactions between pathogens and the cellular surface, our results suggest a parallel mechanism by which sGAGs contribute to pathogen uptake (26). In all sGAG-targeted iBMDMs we discovered a significant decrease in the surface expression of the integrins ITGB2 and ITGAL but no change in the mRNA expression. We propose that one previously overlooked function of sGAGs in pathogen uptake is maintaining the surface expression of critical receptors, including key integrins. How broadly sGAGs control the surface proteome of cells and if this function of sGAGs contributes to other host-pathogen interactions will be of great interest.

Multiple lines of evidence from our study point towards a central role for ITGB2 in the uptake of Mab. Not only was *Itgb2* the number one hit in the genome-wide screen, but we found the top enriched pathway, sGAG-biosynthesis, directly modulates ITGB2 surface expression. ITGB2 is known to complex with different integrin alpha receptors to serve as a key trafficking molecule and complement receptor (49). ITGB2, in complex with ITGAM, was previously shown to modulate LC3-associated phagocytosis (LAP) during *Listeria monocytogenes* infections and control the uptake of *Pseudomonas aeruginosa*, suggesting multiple uptake pathways intersect with ITGB2 (47, 50). The identification of both *Itgam* and *Itgb2* among the top candidates in the screen suggests a key role for this heterodimer in Mab uptake. ITGB2 also complexes with other integrins like ITGAL, yet the contribution of each of these distinct receptors in Mab uptake remains unknown and will be examined in the future (49). While human and murine ITGB2 proteins are highly similar, it will be important to directly test the role of this integrin in Mab uptake in human macrophages (51). In addition, investigating how Mab uptake by distinct receptors alters intracellular trafficking and/or bacterial control will better define key host-pathogen interactions during Mab infections.

How Mab mediates interactions with distinct macrophage uptake pathways remains unknown. Surface glycopeptidolipids (GPLs) are thought to contribute to macrophage infections (52, 53). However, given that both smooth and rough variants of Mab required sGAG-dependent pathways for effective uptake, there must be GPL-independent mechanisms that contribute during infection. One interesting observation from our study was that rough variants are more susceptible to complement-mediated uptake than smooth variants. This suggests a model where GPL evade complement-mediated uptake which may modulate Mab survival. Several other Mab factors have been associated with macrophage infection, including the ATPase EccB4 and glycosyl-diacylated-nondecyl-diols (GDND), but the direct role of these genes in macrophage uptake remain unknown (12, 54-56). More broadly dissecting Mab determinants of macrophage infection will require unbiased bacterial genetic approaches, such as transposon-based approaches that were recently developed in Mab (10, 12). This approach will link macrophage uptake pathways directly with specific Mab-encoded factors.

In addition to understanding how Mab adheres to macrophages, the role of sGAGs during lung infection is important to consider. While Mab is mostly associated with monocytes and macrophages *in vivo*, current mouse models to test Mab pathogenesis remain challenging (55, 57). Sustained Mab infection in mice requires multiple immune deficiencies that disrupt normal host-pathogen interactions (57). In addition, since sGAGs are essential to development, understanding how sGAGs contribute to *in vivo* disease will require the generation of conditional knockout animals in immune knockout backgrounds (58). While improved Mab *in vivo* models are being developed, the approach described here highlights a framework for leveraging unbiased genetic approaches to glean important mechanistic understanding of host-Mab interactions.

Altogether, we optimized a host genetic platform to define important macrophage pathways during Mab infection. By coupling a brightly fluorescent Mab reporter with a genome-wide knockout macrophage library, we are positioned to rapidly screen for diverse phenotypes related to macrophage-Mab interactions. Here, we leveraged this approach to dissect Mab uptake during early infection. Yet, this same approach can be used to understand host pathways that control Mab growth or contribute to macrophage cell death. Thus, our host-focused functional genetic approach will broadly identify potential host-directed therapeutic targets, with the goal of improving treatment success during Mab infection.

## METHODS

### Mab culture conditions, generation of fluorescent reporter, and growth assays

Isogenic pairs of smooth and rough Mab (ATTCC 19977) were used throughout this study. The Mab or *E. coli* mEmerald strains were built by transforming pmV261 hsp60::mEmerald into either the smooth or rough Mab variant by electroporation or DH5α E. coli by heat shock followed by selection on zeocin. All Mab cultures were grown aerobically in Middlebrook 7H9 medium supplemented with 10% Middlebrook OADC (oleic acid, dextrose, catalase, and bovine albumin) at 37°C. mEmerald GFP-expressing strains were propagated in medium containing 5μg/ml zeocin (Invivogen).

To quantify bacterial growth, non-fluorescent and mEmerald GFP-Mab single cell suspensions were inoculated in 7H9 at starting concentration of 5×10^5^ bacteria/ml. To examine antibiotic killing, rifampicin was included at a final concentration of 128 μg/ml. Optical density was quantified by measuring the absorbance at 600nm on a Tecan Spark 20M plate reader at the indicated timepoints.

### Cell culture

J2-virus immortalized murine bone marrow derived macrophages (iBMDMs) and Cas9+ iBMDMs were maintained in Dulbecco’s Modified Eagle Medium (DMEM; Hyclone) supplemented with 10% heat inactivated fetal bovine serum (FBS; Seradigm) as previously described (42). All macrophage cell lines were incubated in 5% CO_2_ at 37°C

### Macrophage infections and uptake assays

Single cell Mab or *E. coli* suspensions were prepared by resuspending logarithmic phase bacteria in DMEM with 10% heat inactivated FBS (Seradigm) followed by a soft spin (800g) to pellet large bacterial clumps. The supernatant was then used to infect macrophages, seeded at 5×10^5^/well in a 12-well plate, at the indicated MOIs for the indicated time points (2-6 hours). Following infection, macrophages were washed with PBS, lifted from plates by scrapping, then fixed in 4% paraformaldehyde. Macrophage uptake was quantified using the BD LSRII flow cytometer (BD Biosciences) at the Michigan State University Flow Cytometry Core. Live and single macrophages were identified using forward and side scatter and the number of infected cells was determined by the fluorescence in the GFP channel. All experiments include an uninfected control to set gates for uptake quantification during analysis that was performed using FlowJo v10. For complement-mediated uptake experiments mEmerald Mab was first incubated in heat-inactivated or active FBS for thirty minutes prior to macrophage infections. For actin polymerization inhibitor experiments, cells were treated with DMSO or Cytochalasin D (Cayman Chemicals) for 2 hours prior to infection.

For the bead uptake assay, 1μm carboxylate-modified polystyrene latex beads expressing yellow-green fluorescence (Sigma-Aldrich) were diluted in PBS to 5×10^8^ beads per ml. Beads were incubated with iBMDMs at the indicated number of latex beads per macrophage for four hours. Following bead exposure, macrophages were washed with PBS and fixed in 4% paraformaldehyde solution. The percent uptake of yellow-green fluorescence latex beads was quantified by flow cytometry as described above.

### CRISPR screen and analysis

Mouse Brie CRISPR knockout pooled library was a gift from David Root and John Doench (Addgene #73633) (42) and infected with our smooth mEmerald-GFP Mab reporter strain at an MOI of five for four hours. Following the infection, macrophages were washed with PBS then fixed with 4% paraformaldehyde. Infected library was then sorted using a BioRad S3e cell sorter to isolate mEmerald+ and mEmerald-macrophages. 1.5-2.5×10^7^ cells were sorted into each bin from triplicate experiments. Genomic DNA was isolated from each sorted population using Qiagen DNeasy kits after reversing DNA crosslinks following a 55°C incubation overnight. Amplification of sgRNAs by PCR was performed as previously described using Illumina compatible primers from IDT (42) and amplicons were sequenced on an Illumina NextSeq 500 at the Genomics Core at Michigan State University.

Sequenced reads were first trimmed to remove any adapter sequence and to adjust for the p5 primer stagger. We used model-based analysis of genome-wide CRISPR-Cas9 knockout (MAGeCK) to map reads to the sgRNA library index without allowing for any mismatch. Subsequent sgRNA counts were median normalized to control sgRNAs in MAGeCK to account for variable sequencing depth. To test for sgRNA and gene enrichment, we used the “test” command in MAGeCK to compare the distribution of sgRNAs in the GFP^+^ and GFP^-^ bins. The MAGeCK output Gene Summary provided a ranked list of genes significantly underrepresented in the mEmerald+ bin based on 4 independent sgRNAs that was used to curate the candidate gene list.

### CRISPR-targeted knockouts

Individual sgRNAs were cloned as previously described using sgOPTI that was a gift from Eric Lander (Addgene plasmid no. 85681) (42). In short, annealed oligos containing the sgRNA targeting sequence were phosphorylated, then cloned into a dephosphorylated and BsmBI (New England Biolabs) digested sgOPTI. To facilitate rapid and efficient generation of sgRNA plasmids with different selectable markers, we used sgOPTI-Blasticidin-Zeocin (BZ) and sgOPTI-Puromycin-Ampicillin (PA) as previously described (42). The simultaneous use of sgOPTI-BZ and sgOPTI-PA plasmids allowed for pooled cloning in which a given sgRNA was ligated into a mixture of BsmBI-digested plasmids. Successful transformants for each plasmid were selected by plating on ampicillin or zeocin in parallel. Next, two sgRNA-plasmid constructs per gene were packaged into lentivirus as previously described (42), then used to transduce Cas9+ iBMDMs. Transductants were selected with blasticidin and/or puromycin then genomic DNA was isolated and PCR was used to amplify edited regions and sent for Sanger Sequencing (Genewiz). The resultant ABI files were used for Tracking of Idels by Decomposition (TIDE) analysis to assess the frequency and size of indels in each population compared to control macrophages (59). Three sgRNAs were targeted per gene and one targeted line was selected for follow-up studies with editing efficiency 60%-95% for each gene. sgRNA sequences and primers for Tide analysis are included in Table S3.

### Quantification of sulfated glycosaminoglycans

Sulfated Glycosaminoglycans (sGAGs) were isolated following the Blyscan Sulfated Glycosaminoglycan Assay (BioVision). In short, macrophages were washed with PBS, then 1ml of Papain Extraction Reagent containing 1mg/ml of Papain (Sigma-Aldrich) was added. Plates were incubated in 5% CO_2_ at 37°C for 1 minute. Cell Suspensions were transferred to 1.5ml microcentrifuge tubes and immediately placed in an ice-water bath. To extract sGAGs, 200μl of cell suspension was transferred to a 1.5ml microcentrifuge tube and incubated at 65°C for three hours.

sGAG were then quantified using the Blyscan Sulfated Glycosaminoglycan Assay Protocol (BioVision). Briefly, 25μl of test sample was transferred into a 1.5ml microcentrifuge tube. Reagent blanks, sGAG standards and test samples were prepared according to the Blyscan Sulfated Glycosaminoglycan Assay General Protocol and adjusted to a 100μl final volume. To stain sGAGs, 1ml of Blyscan Dye Reagent (BioVision) was added to each tube, then tubes were placed in a gentle mechanical shaker for 30 minutes at room temperature. To pellet stained sGAGs, tubes were centrifuged at 13,000xg for 10 minutes, then supernatant was discarded. To release the recovered sGAGs, 500μl of Dissociation Reagent (BioVision) was added to each tube and vortexed until pellets were dissolved. 200μl of reagent blanks, sGAG standards and test samples were transferred to a 96-well plate and absorbance readings at 656nm were obtained from an Agilent BioTek Synergy H1 Hybrid multi-Mode Plate Reader.

### RNA isolation and quantitative real-time PCR

Macrophages were resuspended in 500μL of TRIzol reagent (Life Technologies) and incubated for 5 minutes at room temperature. 100μL of chloroform was added to the homogenate, vortexed and centrifuged at 10,000xg for 18 minutes at 4°C to separate nucleic acids. The clear, RNA containing layer was removed and combined with equal parts ethanol. This mixture was placed into a collection tube and protocols provided by the Zymo Research Direct-zol RNA extraction kit were followed. Quantity and purity of the RNA were checked using a NanoDrop and diluted to 5ng/μL in nuclease-free water.

PCR amplification of the RNA was completed using the One-step Syber Green RT-PCR kit (Qiagen). 25ng of total RNA was added to a master mix reaction of the provided RT mix, Syber green, gene specific primers (5uM of forward and reverse primer), and nuclease-free water. For each biological replicate (triplicate), reactions were conducted in technical triplicates in 96-well plates. PCR product was monitored using the QuantStudio3 (ThermoFisher). The number of cycles needed to reach the threshold of detection (Ct) was determined for all reactions. The mean CT of each experimental sample in triplicate was determined. The average mean of glyceraldehyde 3-phosphate dehydrogenase (GAPDH) was subtracted from the experimental sample mean CT for each gene of interest (dCT).

### Bioinformatics analysis

Gene set enrichment analysis (GSEA) was used to identify enriched pathways in the MAGeCK analyzed dataset based on the full ranked list as done previously. For GSEA analysis, the “GSEA Preranked” function was used to complete functional enrichment using default settings for Kyoto Encyclopedia of Genes and Genomes (KEGG) and Gene Ontology terms. DAVID analysis was used to identify enriched pathways within the positive regulators of Mab uptake. Using the robust rank aggregation (RRA) the top enriched positive regulators (2-fold change with at least 3 sgRNAs) were used as a “candidate list”. Functional analysis and functional annotation analysis were completed, and top enriched pathways and protein families were identified.

### Data availability

Raw sequencing data in FASTQ and processed formats are available for download from NCBI Gene Expression Omnibus (GEO) accession number GSE221413.

### Statistical analysis

Statistical analysis and data visualization were performed using Prism version 9 (GraphPad Software) or Biorender, as indicated in figure legends. Data are presented, unless otherwise indicated, as the mean ± SD and individual data points are shown for each experiment. For parametric data, one-way or two-way ANOVA followed by Tukey’s or Dunnett’s multiple comparisons test were used to identify significant differences between multiple groups.

## Acknowledgments

We would like to acknowledge members of the Olive lab and Robert Abramovitch for helpful discussions and input. We thank the MSU flow cytometry core for their help with instrumentation and analysis as well as the MSU Genomics Core for their help with sequencing. This work was supported by NIH R35GM146795 to AJO, NIFA HATCH 1019371 to AJO and, T32GM142521 to HNG.

